# Positive feedback regulation between RpoS and BosR in the Lyme disease pathogen

**DOI:** 10.1101/2024.09.14.613071

**Authors:** Sajith Raghunandanan, Raj Priya, Gaofeng Lin, Fuad Alanazi, Andrew Zoss, Elise Warren, X. Frank Yang

**Author notes:** Address Correspondence to: Dr. X. Frank Yang Department of Microbiology and Immunology Indiana University School of Medicine Indianapolis, IN 46202.

## Abstract

In *Borrelia burgdorferi*, the Lyme disease pathogen, differential gene expression is primarily controlled by the alternative sigma factor RpoS (σ^S^). Understanding how RpoS levels are regulated is crucial for elucidating how *B. burgdorferi* is maintained throughout its enzootic cycle. Our recent studies have shown that a homolog of Fur/PerR repressor/activator, BosR, functions as an RNA-binding protein that controls the *rpoS* mRNA stability. However, the mechanisms of regulation of BosR, particularly in response to host signals and environmental cues, remain largely unclear. In this study, we revealed a positive feedback loop between RpoS and BosR, where RpoS post-transcriptionally regulates BosR levels. Specifically, mutation or deletion of *rpoS* significantly reduced BosR levels, while artificial induction of *rpoS* resulted in a dose-dependent increase in BosR levels. Notably, RpoS does not affect *bosR* mRNA levels but instead modulates the turnover rate of the BosR protein. Furthermore, we demonstrated that environmental cues do not directly influence *bosR* expression but instead induce *rpoS* transcription and RpoS production, thereby enhancing BosR protein levels. This discovery adds a new layer of complexity to the RpoN-RpoS pathway and suggests the need to re-evaluate the factors and signals previously believed to regulate RpoS levels through BosR.

**IMPORTANCE:** Lyme disease is the most prevalent arthropod-borne infection in the United States. The etiological agent, *Borreliella* (or *Borrelia*) *burgdorferi*, is maintained in nature through an enzootic cycle involving a tick vector and a mammalian host. RpoS, the master regulator of differential gene expression, plays a crucial role in tick transmission and mammalian infection of *B. burgdorferi*. This study reveals a positive feedback loop between RpoS and a Fur/PerR homolog. Elucidating this regulatory network is essential for identifying potential therapeutic targets to disrupt *B. burgdorferi*’s enzootic cycle. The findings also have broader implications for understanding the regulation of RpoS and Fur/PerR family in other bacteria.

## INTRODUCTION

Lyme disease is the most common arthropod-borne infection in the United States, Europe, and Asia (1). The etiological agent, *Borrelia* (or *Borreliella*) *burgdorferi*, perpetuates its life cycle through an enzootic process involving a tick vector and a mammalian host (2). To adapt and survive during this cycle, *B. burgdorferi* undergoes substantial differential gene expression (2–5). Over the past two decades, several regulatory pathways have been identified that govern differential gene expression throughout the enzootic cycle of *B. burgdorferi* (4). Among these, the alternative sigma factor RpoS (σ^S^) has been well recognized as a key regulator, acting as a “gatekeeper” that governs the reciprocal expression of numerous *Borrelia* genes during spirochetal transmission between ticks and mammals (6–8). It activates virulence genes such as *ospC* essential for transmission or infection in vertebrate hosts while suppressing genes such as *ospA* necessary for spirochete survival within the tick vector. Thus, elucidating the molecular mechanism underlying RpoS regulation has become a central focus in *B. burgdorferi* genetics.

Unlike in model organisms such as *Escherichia coli*, RpoS regulation in *B. burgdorferi* is quite unique. The level of RpoS in *B. burgdorferi* is primarily regulated transcriptionally by another alternative sigma factor RpoN (σ^N^), and RpoN and RpoS constitute the RpoN-RpoS (σ^N^– σ^S^) sigma factor cascade or pathway (6, 9, 10). In addition to requiring a bacterial enhancer-binding protein (bEBP) Rrp2 from the σ^N^-type promoter for *rpoS* transcriptional activation (11–15), the *rpoS* expression also requires a Fur/PerR family repressor/activator, BosR (16, 17). Recently, we demonstrated that BosR does not function as a transcriptional regulator as previously proposed to control *rpoS* transcripts but instead, it is a novel RNA-binding protein that directly binds to 5’ untranslated region of the *rpoS* mRNA and controls the turnover rate of the *rpoS* mRNA (18).

*B. burgdorferi* activates the RpoN-RpoS cascade in response to various host and environmental signals, including temperature, cell density, pH, oxygen, carbon dioxide, metals, and short-chain fatty acids (19–28). Although the precise mechanisms by which these signals are integrated into this pathway remain unclear, the prevailing model suggests that host signals and environmental cues regulate RpoS levels through BosR [for review, see (4)]. This hypothesis is primarily based on observations that (1) both the RpoS and BosR levels are influenced by host signals and environmental cues, and (2) BosR governs *rpoS* mRNA levels. However, direct evidence supporting this model is still lacking, and the mechanism by which multiple signals and cellular processes influences RpoS levels remains to be fully elucidated.

In this study, while systematically identifying genes involved in regulating the RpoN-RpoS cascade, we identified two mutants with defects in BosR production. Unexpectedly, both mutants harbored a mutation in the *rpoS* gene, resulting in a truncated RpoS protein. Further investigation revealed that BosR production is regulated by RpoS at the protein level. This finding challenges the current model in which BosR controls RpoS levels. Instead, our results indicate the presence of a novel positive feedback loop between BosR and RpoS. Moreover, we demonstrate that host and environmental factors influence the RpoN-RpoS sigma factor cascade by modulating *rpoS* transcriptional activation, which subsequently affects BosR levels.

## RESULTS

### Truncated RpoS resulted in defective BosR production

To systematically identify genes regulating the RpoN-RpoS cascade, we constructed a transposon library (Tn) in *B. burgdorferi* as previously described (29, 30). Using OspC production as an indicator for RpoS activation, initial screening of Tn mutants by SDS-PAGE analysis identified two transposon mutants that lacked OspC expression (**Fig. 1A**). Sequencing revealed that the transposon was inserted into *bb_0295* and *bb_0421* in Tn-001 and Tn-002, respectively. Western-blotting analyses revealed that Tn-001 and Tn-002 had significantly reduced BosR levels and a loss of RpoS.

**Fig. 1.**
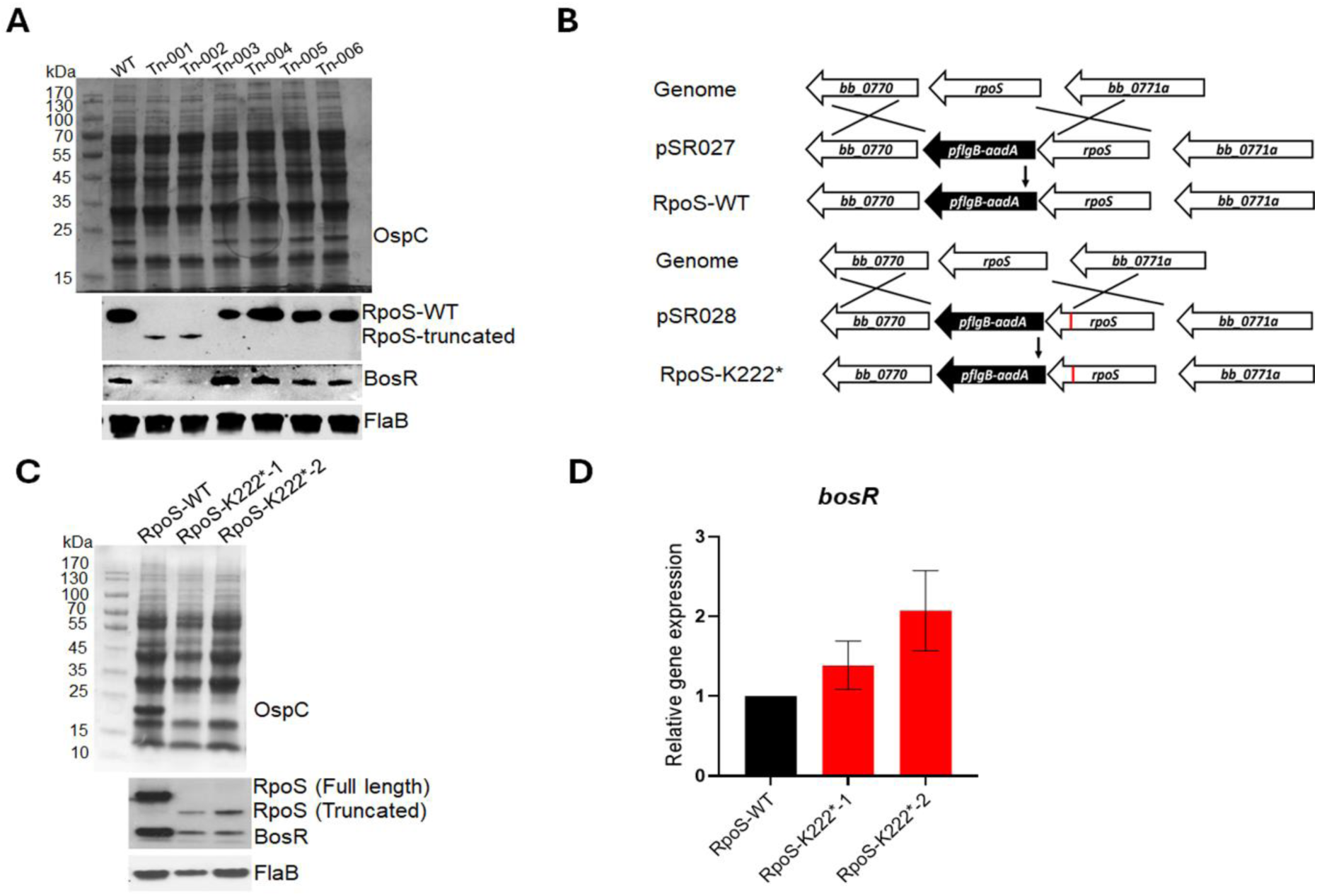
Analyses of *B. burgdorferi* mutants harboring with RpoS truncation. (**A**) Coomassie gel staining and immunoblot analyses of Tn mutants. Wild-type *B. burgdorferi* strain 5A18NP1 and various transposon mutants (labeled at top) were cultured in BSK-II medium at 37°C and harvested at stationary phase. Cell lysates were subjected to SDS-PAGE (top panel) and immunoblot analyses (bottom panel). The bands corresponding to OspC, RpoS, BosR and FlaB are indicated on the right. (**B**) Strategy for constructing a *rpoS* mutant with K222* mutation. pSR027, a suicide vector harboring a wild-type copy of *rpoS* linked to an *aadA* streptomycin-resistant marker; pSR028, a suicide vector identical to pSR027, except harboring a mutated *rpoS* with K222* (depicted in red). pSR027 and pSR028 were transformed into wild-type *B. burgdorferi* strain 5A18NP1, and the resulting strains are designated RpoS-WT and RpoS-K222*, respectively. (**C**) Coomassie gel staining and immunoblot analyses of RpoS-K222* mutants. *B. burgdorferi* strains were cultured and analyzed identical to Fig. 1A. (**D**) Quantitation of *bosR* mRNA levels by qRT-PCR. RNAs were extracted from the cultures in Fig. 1C and subjected to qRT-PCR. The *bosR* mRNA level in strain RpoS-WT were normalized as 1.0. The bars represent the mean values of three independent experiments, and the error bars represent the standard deviation.

Based on the current model, in which BosR controls RpoS, and RpoS subsequently governs OspC production, we initially hypothesized that *bb_0295* and *bb_0421* are important factors that regulate BosR, leading to defective in RpoS and subsequent OspC production. However, we constructed a *bb_0295* and a *bb_0421* null mutant and both mutants showed normal OspC levels (data not shown). Re-constructed Tn mutants by transforming wild-type *B. burgdorferi* with PCR-amplified DNA fragments from Tn-001 and Tn-002 also resulted in normal OspC levels (data not shown), suggesting the OspC defect observed in Tn-001 and Tn-002 was due to additional mutations.

Subsequent genome sequencing revealed a point mutation (T to A) in the *rpoS* ORF in both Tn mutants, which is a nonsense mutation that introduce a stop codon at residue K222, resulting in a 45-amino acid shorter RpoS protein. We hypothesized that the minor, lower molecular weight band detected by anti-RpoS was the truncated RpoS (**Fig. 1A**). Since the region compassing residues 213-263 of RpoS is predicted to be the Helix-Turn-Helix domain critical for DNA-binding, we reasoned that this truncation likely caused the OspC defect. Given that genome sequencing did not reveal additional mutations in Tn-001 and Tn-002, we further hypothesized that RpoS K222* mutation led to impaired BosR levels.

To test this, we constructed a *rpoS* mutant strain with the K222* mutation in the chromosome, designated RpoS-K222* (**Fig. 1B**). The result showed that RpoS-K222* exhibited phenotypes identical to those of Tn-001 and Tn-002: truncated RpoS, abolished OspC production, and notably, reduced BosR levels (**Fig. 1C**). However, no detectable differences were observed in *bosR* mRNA levels in all strains tested (**Fig. 1D**). These findings suggest that RpoS truncation leads to a significant reduction of BosR at the protein level in *B. burgdorferi*.

### RpoS is required for BosR production

The result showing that RpoS truncation by K222* mutation results in decreased BosR protein levels were unexpected, given the well-established notion that BosR controls RpoS levels. To determine whether this phenotype is specific to K222* mutation, we assessed BosR levels in various strains lacking RpoS. As shown in **Fig. 2A**, significant reductions in BosR levels were observed in the *rpoS* deletion mutant, the *rpoN* deletion mutant, and the *rrp2 ^G239C^* mutant. As a control, we included a *bosR* mutant and an *ospC* mutant in the analysis, since we observed that polyclonal anti-BosR antibody often reacts with OspC. However, the anti-BosR antibody used in this study was a monoclonal and specific to BosR. The results from the *bosR* and *ospC* mutants confirmed that the band detected by this anti-BosR monoclonal antibody corresponded to BosR, not OspC (**Fig. 2A**). Additionally, similar to the results shown in **Fig. 1D**, none of the mutants exhibited changes in *bosR* mRNA levels. These findings suggest that the deletion of RpoS leads to a substantial reduction of BosR protein levels.

**Fig. 2.**
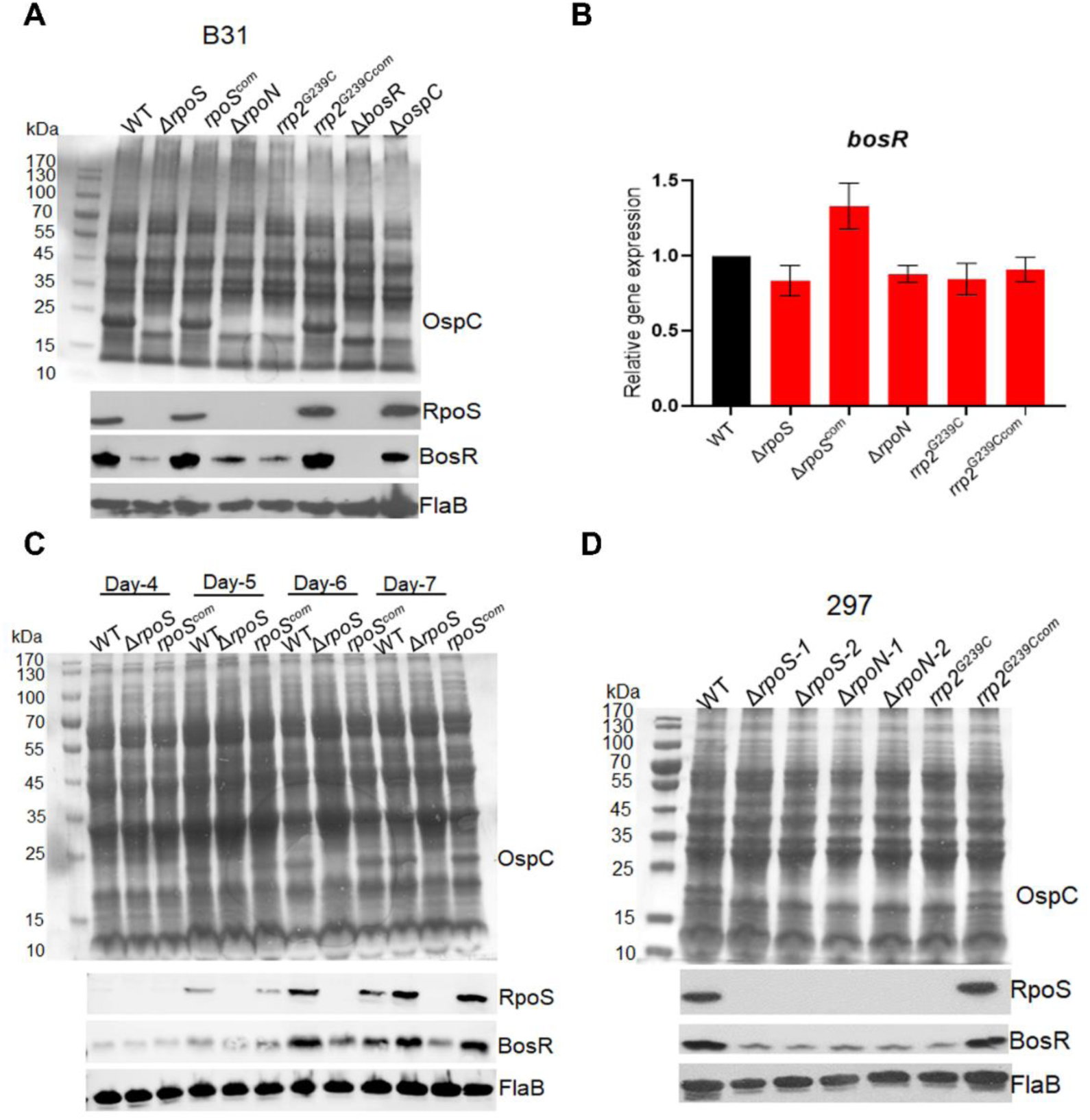
Analyses of BosR levels in various strains lacking RpoS. (**A**) **Coomassie gel staining and Immunoblot analyses.** Wild-type *B. burgdorferi* strain 5A14, the *rpoS* mutant (Δ*rpoS*), *rpoS* complement (*rpoS^com^*), *rpoN* mutant (Δ*rpoN*), *rrp2^G239C^* mutant (*rrp2^G239C^*), *rrp2^G239C^* complement (*rrp2^G239Ccom^*), *bosR* mutant (Δ*bosR*), and *ospC* mutant (Δ*ospC*) were cultured in BSK-II medium at 37°C and harvested at stationary phase (day 6). Cell lysates were subjected to SDS-PAGE (top panel) or immunoblot analyses (bottom panel). The bands corresponding to OspC, RpoS, BosR and FlaB are indicated on the right. (**B**) **Quantitation of *bosR* mRNA levels by qRT-PCR**. RNAs were extracted from the cultures in Fig. 2A and subjected to qRT-PCR. The *bosR* mRNA level in wild-type *B. burgdorferi* 5A14 were normalized as 1.0. The bars represent the mean values of three independent experiments, and the error bars represent the standard deviation. (**C**) **Coomassie gel staining and Immunoblotting of spirochetes harvested at various cell densities.** Wild-type *B. burgdorferi* strain 5A4, *rpoS* mutant, and the complemented strain were cultured in BSK-II medium at 37°C with an initial concentration of 1×10^4^ cells/ml and harvested on day 4, 5, 6 and 7, respectively. Cell lysates were subjected to SDS-PAGE (top panel) or immunoblot analyses (bottom panel). The bands corresponding to OspC, RpoS, BosR and FlaB were indicated on the right. (**D**) **Coomassie gel staining and Immunoblotting of various mutants in 297 background.** Wild-type *B. burgdorferi* strain AH130, *rpoS* mutants (Δ*rpoS*-1 & -2), *rpoN* mutants (Δ*rpoN*-1 & -2), *rrp2^G239C^* mutant (*rrp2^G239C^*), and the *rrp2^G239C^* complemented strain (*rrp2^G239Ccom^*) were cultured were cultured and analyzed identical to Fig. 2A.

The above results were obtained from spirochetes cultured at 37°C and harvested during stationary phase - conditions optimal for RpoS and BosR production. To investigate whether RpoS controlling BosR is specific to the growth phase, wild-type *B. burgdorferi*, the isogenic *rpoS* mutant, and the complemented strains were harvested at various time points (day 4 to day 7). As expected, in both the wild-type and complemented strains, OspC, RpoS, and BosR were induced by increased cell density (concomitantly with decreased culture media pH) (**Fig. 2C**). In the *rpoS* mutant, BosR levels showed a slight increase with increasing in cell density. However, a significant decrease in BosR levels was detected on Days 6 and 7, when RpoS and BosR productions were fully induced in the wild-type type *B. burgdorferi* (**Fig. 2C**). This suggests that RpoS does not influence basal BosR expression but is crucial to induce full BosR production during stationary phase of growth.

To determine whether the regulation of BosR by RpoS is strain-specific, we conducted immunoblot analyses on various mutants lacking RpoS in the strain 297 background. Consistent with the observations in the B31 background, BosR production was significantly decreased in the *rpoS*, *rpoN*, and *rrp2 ^G239C^* mutants (**Fig. 2C**). These findings suggest that the requirement of RpoS for BosR induction at the stationary phase is not strain-specific.

### IPTG-induced *rpoS* expression resulted in a dose-dependent BosR production

To further investigate the control of BosR levels by RpoS, we aimed to determine how artificially varying *rpoS* expression levels would affect BosR protein levels. To this end, we employed a previously constructed shuttle vector containing the *rpoS* ORF along with 50 bp of *rpoS* 5’UTR driven by a *lac* promoter (designated *lacp*-UTR*_rpoS_*-*rpoS*, **Fig. 3A**) (18). This plasmid was then transformed into a *rpoS* mutant. As expected, IPTG induction in the *rpoS* mutant carrying the *lacp*-UTR*_rpoS_*-*rpoS* led to an increase in RpoS levels in a dose-dependent manner (**Fig. 3B**). In the uninduced culture, low or basal levels of BosR protein was detected in the *rpoS* mutant, but BosR levels increased following IPTG induction (50 to 125 µM), correlating with the increase in RpoS levels (**Fig. 3B**). However, IPTG-induced *rpoS* expression did not affect *bosR* mRNA levels (**Fig. 3C**).

**Fig. 3.**
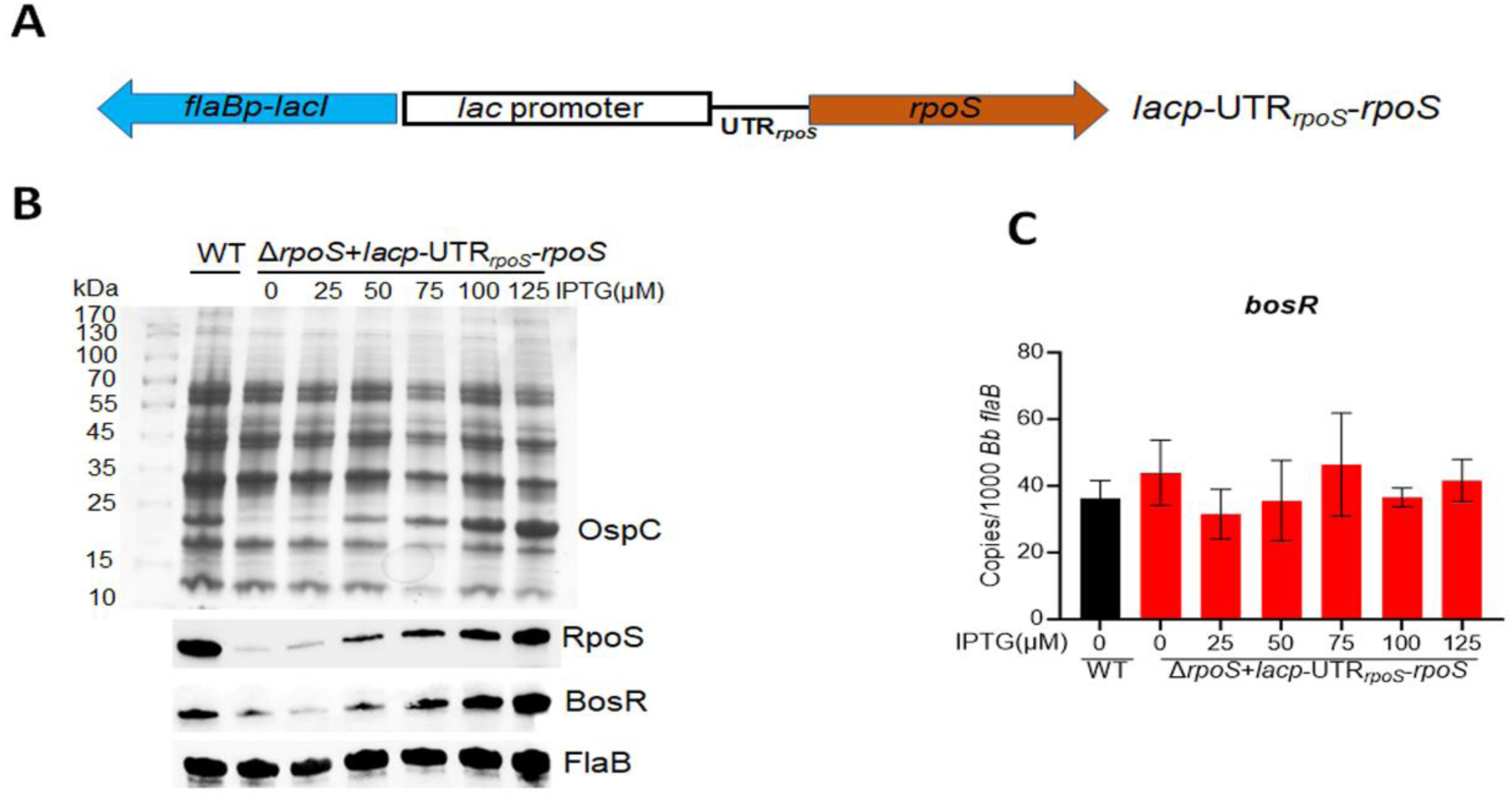
Influence of BosR levels by IPTG-induced *rpoS* expression. (**A**) Schematic representation of the shuttle vector carrying an IPTG-inducible *rpoS* gene (*lacp*-UTR*_rpoS_*-*rpoS*). The blue arrow labeled as *flaBp-lacI* is a *flaB* promoter-driven *lacI* gene. The *lac* promoter is fused with a fragment containing 5’UTR*_rpoS_* and *rpoS* ORF (brown arrow). (**B**) **Coomassie gel staining and Immunoblot analyses**. wWld-type *B. burgdorferi* strain 5A14 and the isogenic *rpoS* mutant harboring *lacp*-UTR*_rpoS_*-*rpoS* were cultured in BSK-II medium with an initial concentration of 1x10^4^ spirochetes/ml and with various concentrations of IPTG (indicated on top). Spirochetes were harvested on day 6 and subjected to SDS analysis (top panel) or immunoblotting (bottom panel) using monoclonal antibodies against RpoS, BosR or FlaB (loading control). (**C**) Quantitation of *bosR* mRNA levels by qRT-PCR. RNAs were extracted from the cultures in (**B**) and subjected to qRT-PCR. The values represent the *bosR* mRNA copies normalized to 1000 copies of *B. burgdorferi flaB* mRNA. The bars represent the mean values of three independent experiments, and the error bars represent the standard deviation.

We recently reported that BosR binds to *rpoS* 5’UTR region and governs the turnover rate of *rpoS* mRNA (18). Thus, the stability of *rpoS* mRNA transcribed from the *lacp*-UTR*_rpoS_*-*rpoS* construct used above is influenced by BosR, complicating the interpretation of the result. Additionally, binding to the RNA target often influences the stability of bacterial RNA-binding proteins (31). To simplify the study, we employed two additional IPTG-inducible *rpoS* shuttle vectors: *lacp*-UTR*_lac_*-*rpoS*, in which the *rpoS* 5’UTR is deleted (retaining only the 5’UTR from the *lac* promoter), and the shuttle vector *lacp*-UTR*_flaB_*-*rpoS*, where the *rpoS* 5’UTR is replaced with the *flaB* 5’UTR (**Fig. 4A**) (18). We previously demonstrated that the *rpoS* mRNA transcribed from both constructs no longer requires BosR for stability, allowing for BosR-independent *rpoS* expression (18). Accordingly, an *rpoS* mutant harboring either *lacp*-UTR*_lac_*-*rpoS* or *lacp*-UTR*_flaB_*-*rpoS* were subjected to immunoblot analysis to assess BosR levels. As expected, IPTG induction in the *rpoS* mutant carrying either of these plasmids resulted in increased RpoS production in a dose-dependent manner (**Fig. 4B & 4C**). Consistent with the pattern observed in **Fig. 3B**, BosR exhibited a similar dose-dependent increase in response to IPTG-induced RpoS production (**Fig. 4B & 4C**). These results collectively support that notation that RpoS regulates BosR protein levels in *B. burgdorferi*.

**Fig. 4.**
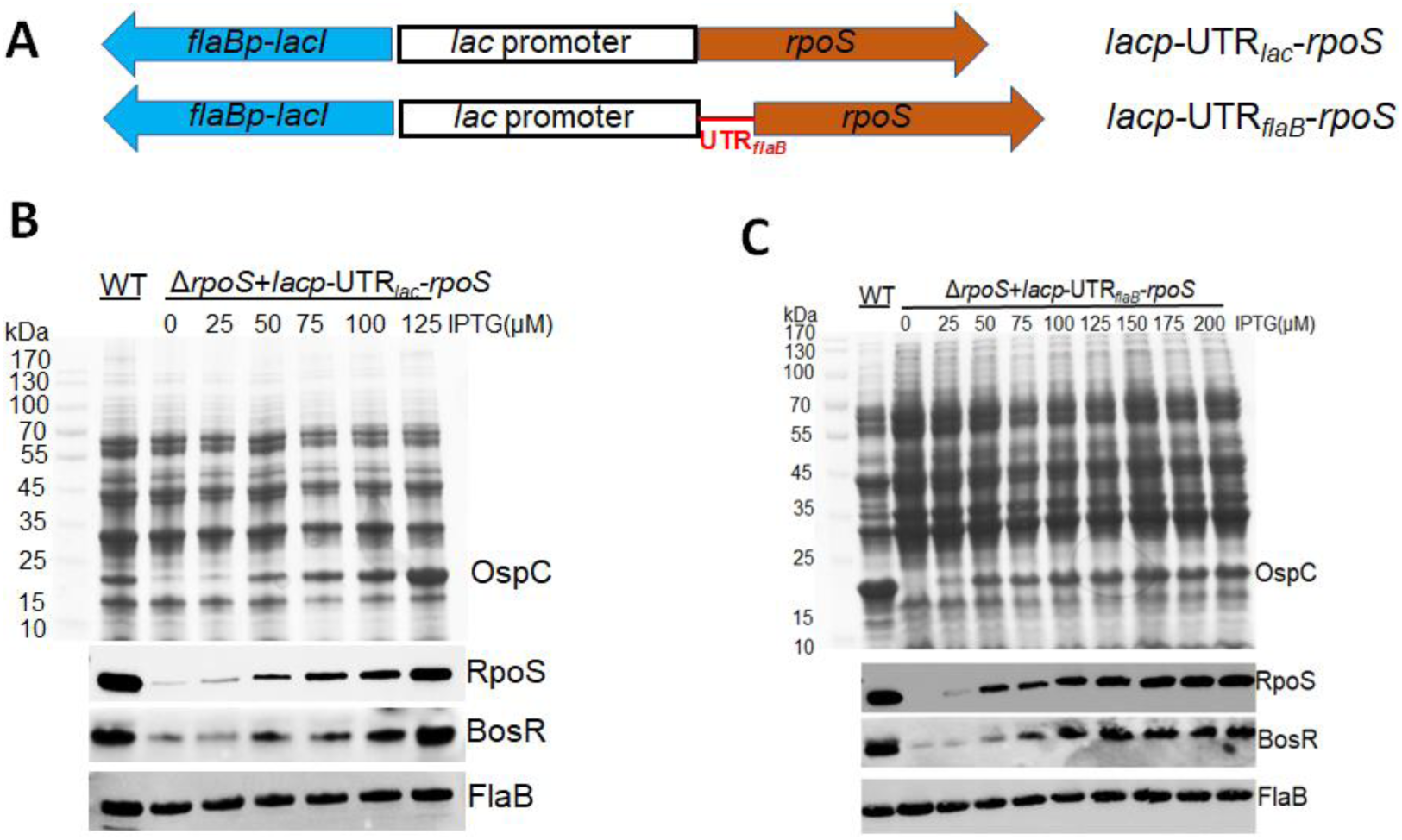
Influence of BosR levels by IPTG-induced, BosR-independent *rpoS* expression. (**A**) Schematic representation of the shuttle vectors *lacp*-UTR*_lac_*-*rpoS* and *lacp*-UTR*_flaB_*-*rpoS*. The blue arrow labeled as *flaBp-lacI* is a *flaB* promoter-driven *lacI* gene. *lacp*-UTR*_lac_*-*rpoS*, the *lac* promoter is fused with the *rpoS* gene (brown arrow) in which the *rpoS* 5’UTR is deleted (retaining only the 5’UTR from the *lac* promoter); *lacp*-UTR*_flaB_*-*rpoS*, the *lac* promoter is fused with the *rpoS* gene where the *rpoS* 5’UTR is replaced with the *flaB* 5’UTR (highlighted in red). (**B & C**) **Coomassie gel staining and Immunoblot analyses**. Wild-type *B. burgdorferi* strain 5A14 and the isogenic *rpoS* mutant harboring *lacp*-UTR*_lac_*-*rpoS* (**B**) or *lacp*-UTR*_flaB_*-*rpoS* (**C**) were cultured in BSK-II medium with an initial concentration of 1 x 10^4^ spirochetes/ml and with various concentrations of IPTG (indicated on top). Spirochetes were harvested on day 6 and subjected to SDS analysis (top panel) or immunoblotting (bottom panel) using monoclonal antibodies against RpoS, BosR or FlaB (loading control).

### IPTG-induced *bosR* expression requires RpoS for BosR production

The result above suggests that RpoS does not influence *bosR* transcription. To gather further evidence that RpoS regulates BosR production at the protein level, rather than the transcription level, we investigated the effect of RpoS on BosR levels produced from IPTG-induced *bosR* transcription. Thus, a shuttle vector harboring an IPTG-inducible *bosR* ORF (designates *lacp*-UTR*_lac_*-*bosR*, **Fig. 5A**), was transformed into the *bosR*, *rpoS*, and *rpoN* deletion mutants, respectively. As expected, IPTG induction in the *bosR* mutant carrying *lacp*-UTR*_lac_*- *bosR* resulted in a dose-dependent increase in BosR protein levels (**Fig. 5B**). In contrast, no increase in BosR protein levels was detected in the *rpoS* or *rpoN* mutants upon IPTG induction (**Fig. 5B**), despite an observed increase in *bosR* mRNA in both mutants in response to IPTG (**Fig. 5C**). These findings indicate that IPTG-induced *bosR* expression requires RpoS for the full production of BosR protein.

**Fig. 5.**
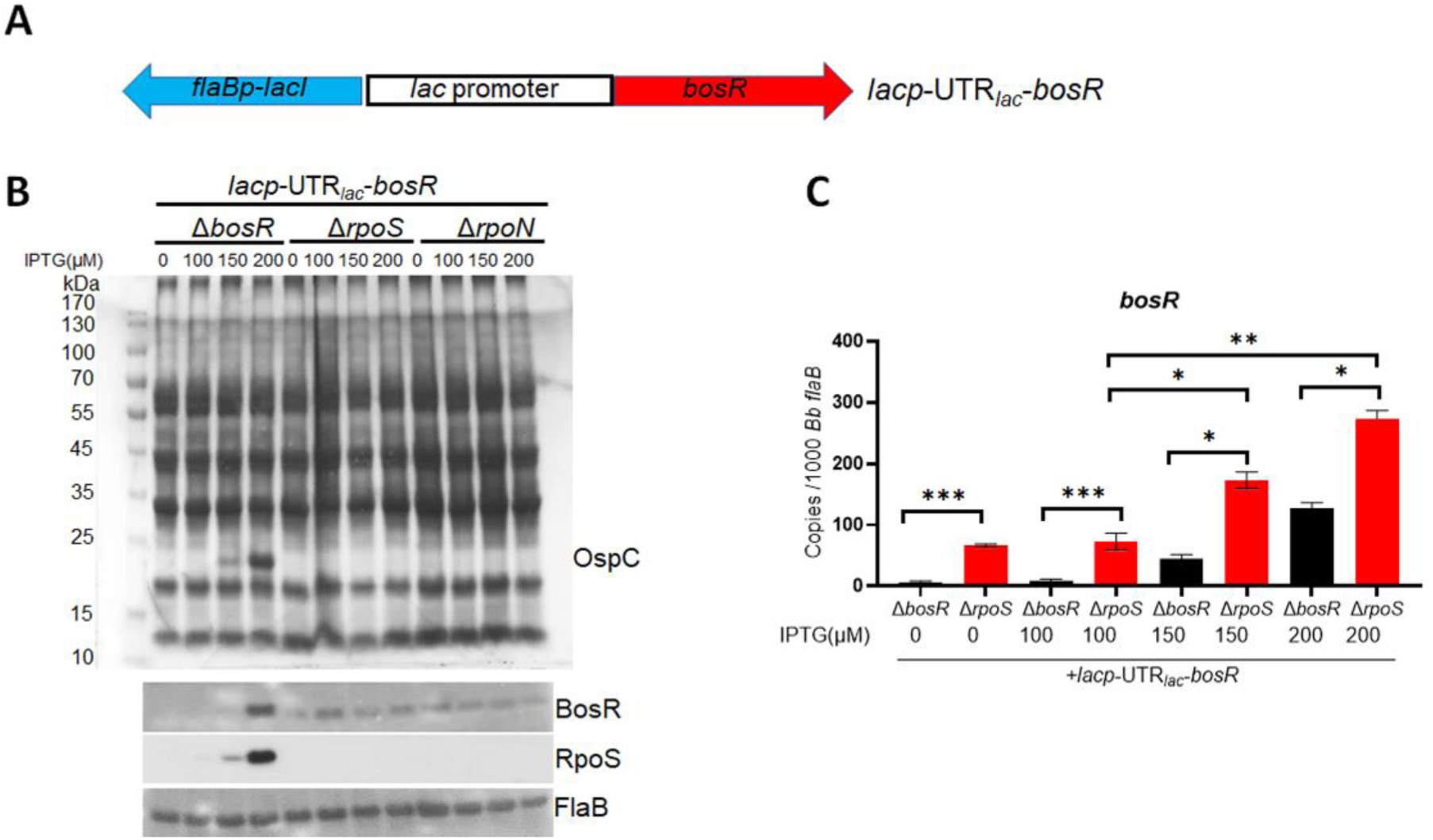
Influence of RpoS on BosR protein levels produced from IPTG-induced *bosR* transcription. (**A**) Schematic representation of the shuttle vector carrying an IPTG-inducible *bosR* gene (*lacp*-UTR*_lac_*-*bosR*). The blue arrow labeled as *flaBp-lacI* is a *flaB* promoter-driven *lacI* gene. The *lac* promoter is fused with the *bosR* ORF (red arrow, which has a 5’UTR within the lac promoter sequence, UTR*_lac_*). (**B**) **Coomassie gel staining and Immunoblot analyses**. The *bosR, rpoN* and *rpoS* mutants harboring *lacp*-UTR*_lac_*-*bosR* plasmid were cultured in BSK-II medium with various concentrations of IPTG (indicated on bottom). Spirochetes were harvested at stationary phase and then subjected to SDS analysis (top panel) or immunoblotting (bottom panel). (**C**) Quantitation of *bosR* mRNA levels by qRT-PCR. RNAs were extracted from (**B**) and subjected to qRT-PCR analyses. The values represent the *bosR* mRNA copies normalized to 1000 copies of *B. burgdorferi flaB* mRNA. The bars represent the mean values of three independent experiments, and the error bars represent the standard deviation. ***, *p* < 0.0001, **, *p* < 0.001, *, *p* < 0.01, using one-way ANOVA.

### RpoS controls BosR protein levels in mammalian host-adapted spirochetes

Spirochetes grown at 37°C in stationary phase activate the RpoN-RpoS pathway, providing a valuable model for investigating the regulatory mechanism of this pathway. However, these conditions do not fully capture the extent of RpoS activation observed during mammalian infection, such as very high level of OspC production and diminished OspA production characteristic of spirochetes in this environment (8, 25). To determine the influence of RpoS on BosR levels under conditions that mimic the mammalian host environment, we cultivated spirochetes using a dialysis membrane chamber (DMC) implanted in the peritoneal cavities of rats (8, 32). As shown in **Fig. 6**, wild-type and the *rpoS*-complemented strains grown under DMC conditions produced high levels of OspC and undetectable levels of OspA, consistent with the expected host-adapted phenotype. The *rpoS* mutant, as previously reported, failed to activate OspC and repress OspA (8). Notably, BosR was virtually undetectable in the *rpoS* mutant under DMC conditions (**Fig. 6**), implying an essential role of RpoS in BosR production during mammalian infection.

**Fig. 6.**
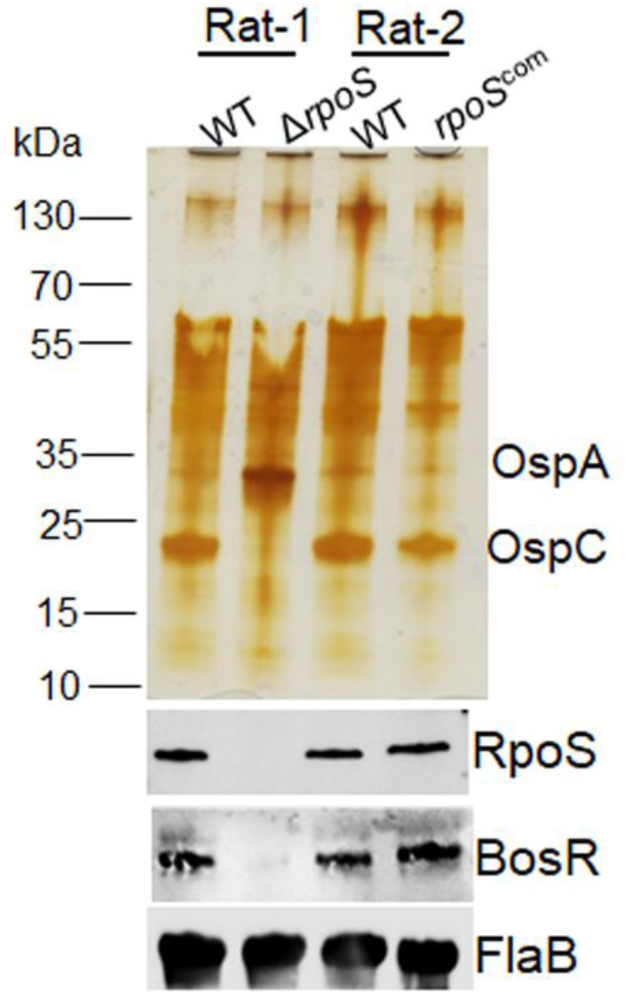
Influence of RpoS on BosR levels under mammalian host-adapted conditions. Wild-type *B. burgdorferi* strain 5A4 (WT), the isogenic *rpoS* mutant (Δ*rpoS*) and the complementation stain (*rpoS*^com^) were cultivated in DMCs. Thirteen days after implantation, spirochetes were harvested and subjected to silver staining and immunoblot analyses. Bands corresponding to OspA, OspC, RpoS, BosR and FlaB (loading control) are indicated on the left.

### RpoS regulates BosR protein turnover rate

To explore the mechanism by which RpoS regulates BosR protein levels, we examined the impact of RpoS on the turnover rate of BosR protein, a common bacterial mechanism for controlling protein levels (33). Accordingly, we compared the turnover rates of BosR protein among wild-type *B. burgdorferi*, the *rpoS* mutant, and the *rpoS*-complemented strains, by treating the cultures with spectinomycin, a bacterial protein translation inhibitor (34, 35). In both the wild-type and *rpoS*-complemented spirochetes, BosR protein levels remained relatively stable during 24 hrs of spectinomycin treatment (**Fig. 7**). In contrast, in the *rpoS* mutant, BosR protein levels began to decrease 6 hrs after spectinomycin treatment, and were completely diminished after 12 and 24 hrs, despite the low basal level of BosR protein present before the treatment compared to those in the wild-type and *rpoS*-complemented spirochetes (**Fig. 7**). FlaB levels in the *rpoS* mutant were not altered by spectinomycin treatment. These findings suggest that RpoS controls the turnover rate of BosR protein.

**Fig. 7.**
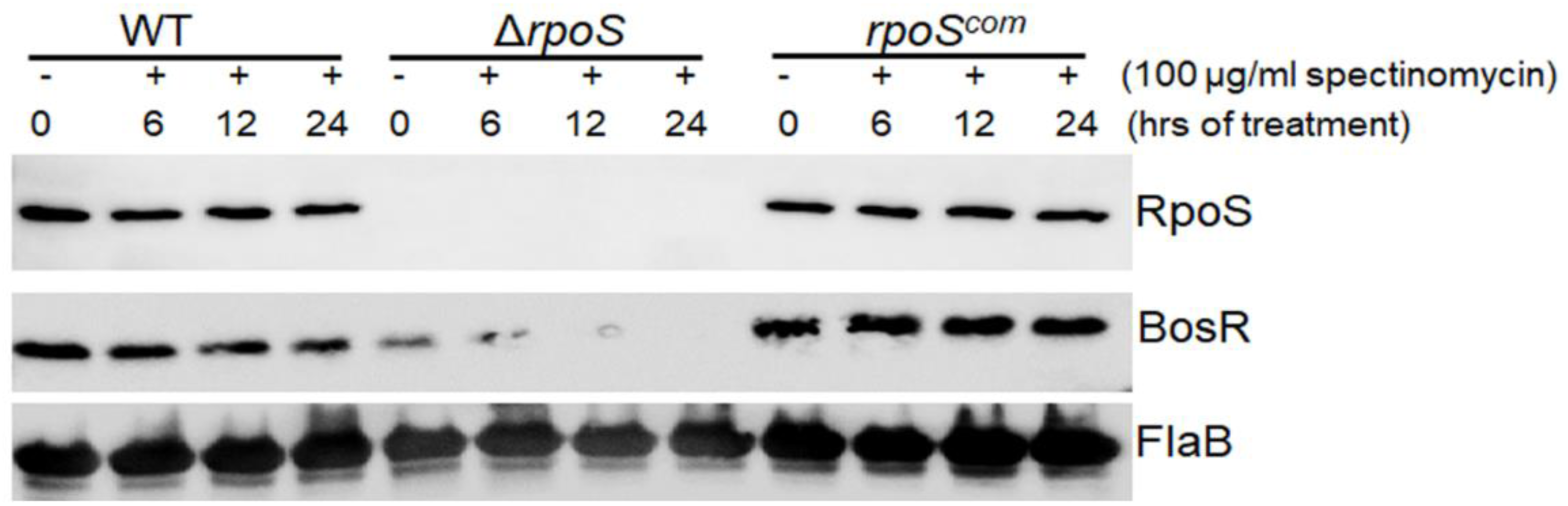
BosR protein turnover assay. Wild-type *B. burgdorferi* strain 5A14, *rpoS* mutant (Δ*rpoS*), or *rpoS* complement (*rpoS^com^*) spirochetes were cultured in BSK-II medium at 37°C. On day 7 during stationary phase of growth, translational arrest was performed by treating the cultures with spectinomycin. Spirochetes were harvested at various time points as indicated and subjected to immunoblotting. Experiments were repeated in three independent biological replicates. A representative image is shown here.

### Environmental cues regulate *rpoS* transcription

The current model suggests that environmental cues induce RpoS production through BosR, based on BosR’s role in governing *rpoS* expression. However, our findings indicate that RpoS also modulates BosR protein levels, necessitating a re-examination of how environmental cues induce RpoS production. To investigate this, we first assessed the impact of environmental cues on the promoter activity of the *bosR* or *rpoS* gene.

Wild-type *B. burgdorferi* was transformed with a shuttle vector containing a luciferase reporter (*luc* ORF) driven by the *bosR* promoter (pOY463, which includes 2.1 kb upstream of the BosR ORF and contains both the P1 and P2 *bosR* promoters) (36) (**Fig. 8A Top**). Another set of spirochetes was transformed with a shuttle plasmid containing a luciferase reporter driven by the σ^54^-type *rpoS* promoter (pJK002, which includes 75 bp upstream of the *rpoS* ORF and lacks putative BS1 and BS2 BosR binding sites) (**Fig. 8B Top**). The constructed *B. burgdorferi* strains were grown either at 23°C or 37°C and harvested at mid-logarithmic or stationary phase. RNAs were then extracted and subjected to qRT-PCR analyses to assess the levels of the *luc* reporter RNA, as well as native *rpoS* and *bosR* RNA.

**Fig. 8.**
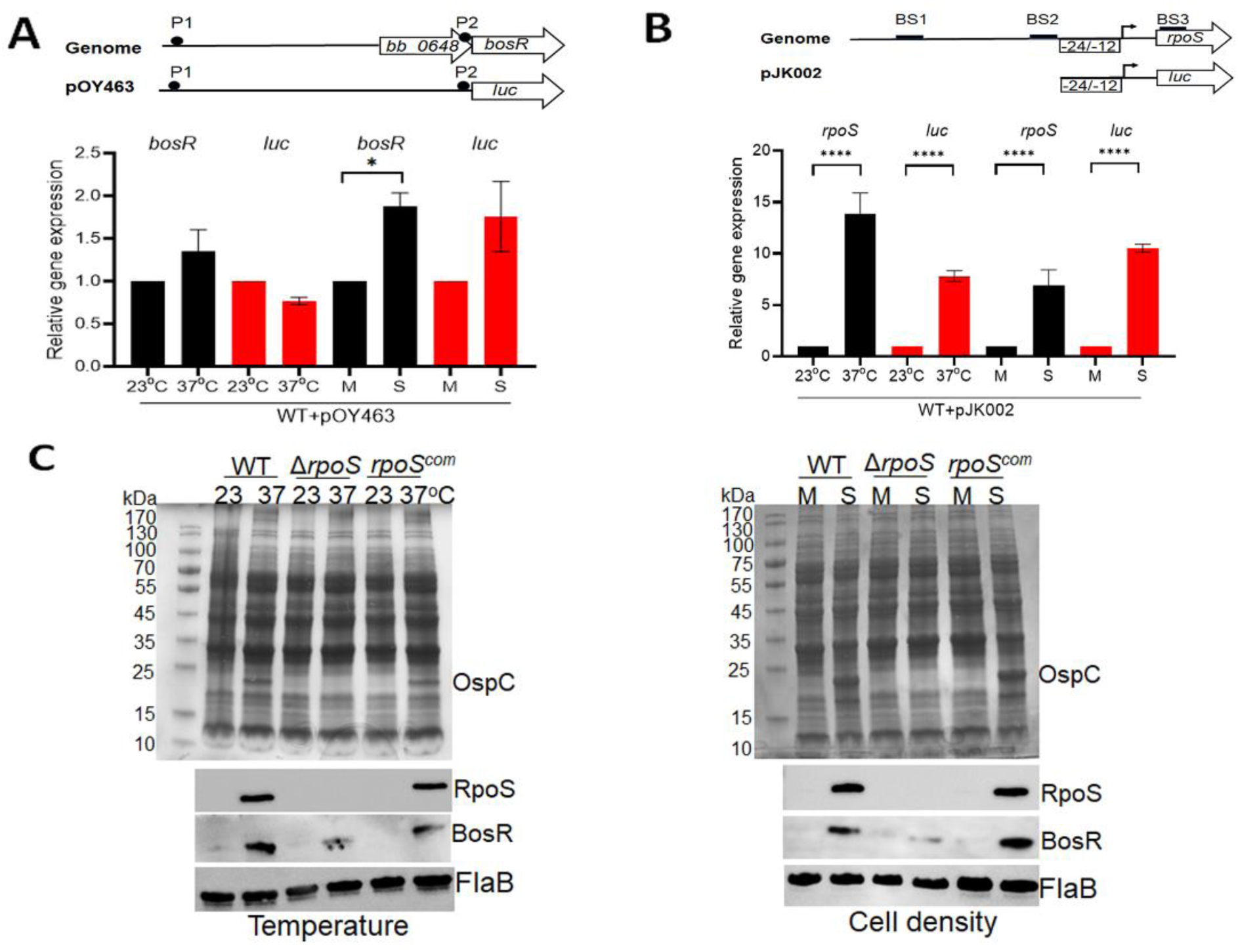
Effects of temperature and cell density on transcriptional activation of *rpoS* and *bosR*. (**A**) Analyses of the luciferase reporter driven by the *bosR* promoter. Top panel shows a schematic representation of the reporter construct. The upper diagram illustrates the organization of *bosR* gene in the genome. The lower diagram depicts the *luc* gene fused to the full length of the *bosR* promoter in shuttle vector pOY463. The putative *bosR* promoter P1 and P2 are highlighted in black circles. For qRT-PCR analyses (Bottom panel), wild-type *B. burgdorferi* strain B31 carrying pOY463 (WT+pOY463) was cultured in BSK-II medium either at 23°C and 37°C and harvested at mid-log (M) or stationary (S) phases. RNAs were extracted and subjected to qRT-PCR analyses. The expression levels of both *bosR* and *luc* isolated from 23°C and mid-log culture were set as 1.0. (**B**) Analyses of the luciferase reporter driven by the *rpoS* promoter. The upper diagram illustrates the organization of *rpoS* gene in the genome. The lower diagram depicts the *luc* gene fused to the sigma54-type minimal *rpoS* promoter in shuttle vector pJK002. For qRT-PCR analyses (Bottom panel), spirochetes were cultured in BSK-II medium either at 23°C and 37°C and harvested at mid-log (M) or stationary (S) phases. RNAs were extracted and subjected to qRT-PCR analyses. The expression levels of both *rpoS* and *luc* isolated from 23°C and mid-log culture were set as 1.0. All bars represent the mean values of three independent experiments, and the error bars represent the standard deviation. \*\*\*\**p* < 0.0001 respectively using one-way ANOVA. (**C**) **Coomassie gel staining and immunoblot analyses**. Wild-type *B. burgdorferi* strain 5A4, the isogenic *rpoS* mutant (Δ*rpoS*), the *rpoS*- complemented strain (*rpoS^com^*), were cultured in BSK-II medium at 23°C and 37°C (**Left**) and harvested at mid-log phase (day 2) and stationary phase (day 6) (**Right,** 37°C). Cell lysates were subjected to SDS-PAGE (top panel) or immunoblot analyses (bottom panel). The bands corresponding to OspC, RpoS, BosR and FlaB were indicated on the right.

In wild-type *B. burgdorferi* harboring pOY463, no significant changes in *luc* or bosR mRNA levels were observed when comparing spirochetes grown at 23 vs 37°C, suggesting that elevated temperature does not induce *bosR* promoter activity (**Fig. 8A, Bottom**). However, a 1.8- fold increase in *bosR* mRNA levels was observed when comparing mid-log to stationary phase cultures (**Fig. 8A, Bottom**). Although stationary phase growth appeared to increase *luc* mRNA levels by approximately 1.7-fold, this increase was not statistically significant. Nevertheless, the moderate increases in *bosR* and *luc* mRNA levels do not fully account for the substantial induction of BosR observed under stationary phase conditions (**Fig. 2C**). In contrast, both *rpoS* and *luc* mRNA levels increased to 7-14 folds in response to elevated temperature and increased cell density in wild-type *B. burgdorferi* harboring pJK002 (**Fig. 8B, Bottom**). These findings suggest that environmental cues induce the activation of *rpoS* transcription.

We then investigated whether environmental cues-induced BosR production is RpoS- dependent. Wild-type *B. burgdorferi*, the isogenic *rpoS* mutant, and the *rpoS*-complemented strain were grown at 23 or 37°C and harvested at mid-log or stationary phase. Results showed that *rpoS* deletion significantly reduced temperature- and cell density-induced BosR production (**Fig. 8C)**. These combined findings suggest that environmental cues induce *rpoS* transcription and RpoS production, which in turn promotes BosR protein levels.

## DISCUSSION

RpoS serves as a master regulator that orchestrates the differential expression of numerous genes during the enzootic cycle of *Borrelia burgdorferi*. Given its critical and complex role, spirochetes have evolved multiple mechanisms to modulate RpoS levels in response to various host and environmental signals at different stages of the cycle. The current model proposes that environmental signals and *Borrelia* factors modulate RpoS levels through BosR, based on the findings that BosR governs *rpoS* expression and its own levels are influenced by these signals and factors. In this study, however, we present a novel positive feedback loop between RpoS and BosR. Our findings show that RpoS post-transcriptionally regulates BosR levels, while environmental cues stimulate *rpoS* transcription and RpoS production, thereby enhancing BosR protein levels. These findings not only introduce a new layer of regulation to the existing paradigm of the RpoN-RpoS cascade, but also call for a re-evaluation of all factors and signals previously believed to primarily influence RpoS levels through BosR.

Several lines of evidence support the conclusion that RpoS regulates BosR protein levels: (1) Mutants deficient in RpoS production exhibited reduced BosR protein levels despite unchanged *bosR* mRNA levels (**Fig. 1 & 2**); (2) IPTG-induced RpoS production led to a dose-dependent increase in BosR protein levels (**Fig. 3 & 4**); (3) IPTG-induced *bosR* expression required RpoS for BosR protein production (**Fig. 5**); (4) Both environmental cues and DMC conditions required the presence of RpoS to induce BosR production (**Fig. 6 & 8**). We further demonstrate that RpoS modulates BosR levels by influencing protein turnover rate (**Fig. 7**), though the precise mechanism remains to be elucidated. Given that BosR is a newly identified RNA-binding protein, one plausible mechanism is that RpoS could regulate BosR by regulating the availability of BosR’s RNA binding targets, as bacterial RNA-binding proteins are often stabilized through their interactions with RNA (31). Thus far, the only RNA-binding site identified for BosR is the 5’UTR region of *rpoS* (18). However, the regulation of BosR protein levels by RpoS does not appear to involve BosR binding to *rpoS* RNA, as IPTG-induced RpoS production from *rpoS* mRNA, with or without the 5’UTR binding site for BosR, yielded similar results (**Fig. 4B & 4C**). It remains possible that RpoS facilitates BosR binding to other, as yet unidentified, RNA targets, thereby stabilizing BosR. Nevertheless, regulation of BosR By RpoS is likely indirect, as RpoS, being a global regulator, controls the expression of numerous genes in *B. burgdorferi*. RpoS may influence expression of factors within proteolysis pathways involved in RosR degradation, a common regulatory mechanism of bacterial protein turnover regulation (37).

The notion that the induction of BosR levels by environmental cues and other factors mainly occurs at protein levels rather than at mRNA levels has been reported previously by several groups (23, 24, 36, 38–40). In ticks and mammals, regulation of *bosR* expression levels has been observed (36, 40, 41), but determining whether regulation at the protein level *in vivo* is challenging and remain to be determined. Although our results demonstrate the pivotal role of regulation of BosR protein levels by RpoS, it does not diminish the importance of regulation of *bosR* at the transcriptional level. Elegant work done by Ouyang et al., showed that the transcription of *bosR* in *B. burgdorferi* is chiefly governed by a σ^70^-type promoter and BosR can auto-regulate its own expression at this promoter (36).

In addition to RpoS-dependent BosR production, there is also RpoS-independent BosR production, as a basal level of BosR protein was observed in RpoS-deficient strains (**Fig. 2C**). This basal level was detected even during early- and mid-log phase cultures when RpoS was absent (**Fig. 2C**). In contrast, RpoS-dependent BosR production was most prominent during the stationary phase of growth. As such, this “RpoS-dependent BosR production” phenomenon can be overlooked if the culture conditions are not optimal for high RpoS levels (as indicated by a prominent OspC band in the Coomassie-stained gel). On the other hand, while RpoS-dependent BosR production is critical for high levels of BosR production, a constitutive RpoS-independent, basal level of BosR production is likely essential for initial production of RpoS and for initiation of the positive feedback loop between RpoS and BosR. During the shift from uninduced to induced conditions, *rpoS* transcription is activated. The newly synthesized *rpoS* mRNA requires the presence of BosR to bind to *rpoS* 5’UTR, preventing *rpoS* mRNA degradation and enabling RpoS production. The produced RpoS subsequently enhances BosR protein stability, leading to increased BosR protein levels which allow further accumulation of *rpoS* mRNA and higher RpoS production (**Fig. 9**). Thus, this positive feedback loop between RpoS and BosR enables rapid amplification of RpoS production in response to environmental changes during nymphal tick feeding.

**Fig. 9.**
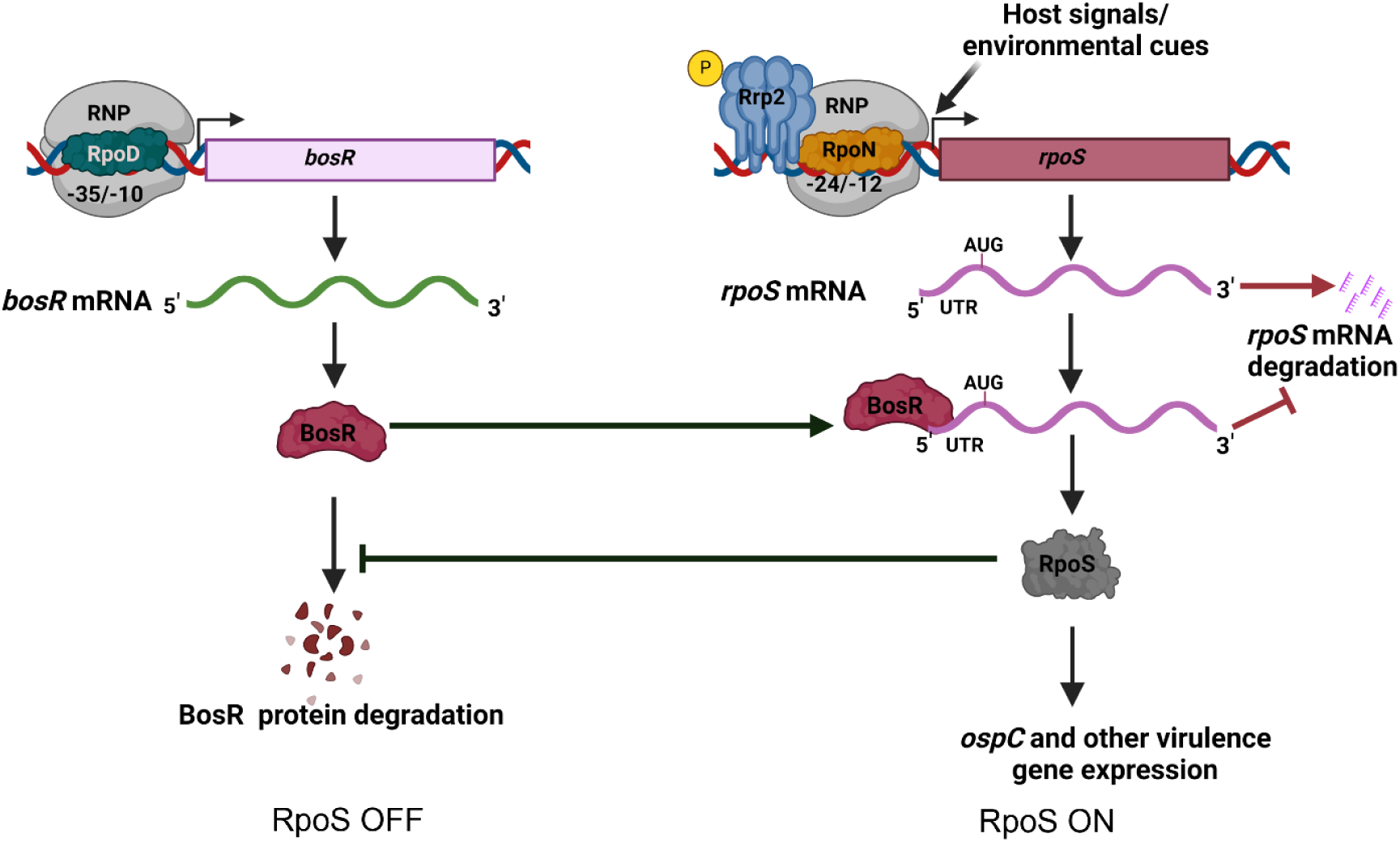
Proposed model of the positive feedback loop between RpoS and BosR. During the RpoS OFF stages of the *B. burgdorferi* enzootic cycle, BosR is produced at a basal level. In the RpoS ON stages, host and environmental signals activate the *rpoS* transcription via an unknown mechanism, newly transcribed *rpoS* mRNA is protected from degradation by the basal level of BosR binding to the *rpoS* 5’UTR region. The produced RpoS protein then inhibits the degradation of BosR, leading to an increase in BosR protein levels, which in turn protects more *rpoS* mRNA from degradation, resulting in a rapid increase in RpoS levels.

Feedback regulation of alternative sigma factor RpoS has been reported in several bacterial species. In *Vibrio cholerae*, cellular levels of RpoS during energy-deprived growth conditions represses the transcription of the response regulator gene *rssB* (42), which is crucial for the proteolytic degradation of RpoS by the ClpXP protease complex (43, 44). During normal growth conditions, elevated RssB levels directly lead to RpoS protein degradation. This feedback regulation between RpoS and RssB controls the motility and colonization in *V. cholerae* (45). Another example is the negative feedback loop between RpoS and anti-adaptor IraP in *E. coli* in response to various stresses (46). However, feedback regulation of RpoS with the involvement of Fur/PerR family proteins has not been observed in other bacteria.

In summary, this study uncovered a positive feedback loop between RpoS and BosR, revealing that not only does BosR regulate RpoS levels, but RpoS also regulates BosR levels (**Fig. 9**). Furthermore, host signals (DMC) and environmental cues primarily stimulate RpoS production by inducing *rpoS* transcription, contrary to previous notion that these signals increase BosR levels which in turn, increase RpoS production. These findings raise several interesting questions: (1) If not through BosR, how do environmental signals regulate *rpoS* transcriptional activation? Besides BosR, Rrp2 and RpoN are two key regulators for *rpoS* transcription, but their levels remain relatively stable across different growth conditions, and Rrp2 phosphorylation is constitutively on in replicating spirochetes as it is crucial for *B. burgdorferi* viability (11, 15, 47, 48). Therefore, the precise mechanism by which environmental signals trigger *rpoS* transcription remains to be elucidated. (2) Several factors, including but not limited to BadR, ppGpp, Rrp1, PlzA, and BmtA, have been identified to regulate RpoS by modulating BosR levels (24, 38, 49–53). Since these conclusions were largely based on the assumption that BosR controls RpoS, could these factors, in fact, control RpoS first, subsequently influencing BosR levels? (3) Conflicting results exist regarding how *ospA* expression is repressed by the RpoN-RpoS pathway. Some studies suggest that BosR directly binds to the *ospA* promoter and represses *ospA* expression (54, 55), while others propose that since the loss of RpoS abolishes *ospA* repression, RpoS, not BosR, is responsible (53, 56). The new insight of RpoS regulating BosR presented in this study may help resolve these conflicting findings.

## MATERIALS AND METHODS

### *B. burgdorferi* strains and culture conditions

Low-passage, virulent *B. burgdorferi* strain 5A18NP1, AH130 and 5A14 were kindly provided by Drs. H. Kawabata and S. Norris, University of Texas Health Science Center at Houston. Spirochetes were cultivated in Barbour-Stoenner-Kelly (BSK-II) medium supplemented with 6% normal rabbit serum (Pel-Freez Biologicals, Rogers, AR) at 37°C with 5% CO2 (57). Appropriate antibiotics were added to the cultures at the time of growth, with final concentrations of 300 μg/ml for kanamycin, 50 μg/ml for streptomycin, and 50 μg/ml for gentamicin, respectively. All the constructed plasmids were maintained in *Escherichia coli* strain DH5α. The antibiotic concentrations used for *E. coli* selection were as follows: streptomycin, 50 μg/ml; gentamicin, 15 μg/ml and rifampicin, 50 μg/ml, respectively. The *B. burgdorferi* strains and plasmids used in this study are listed in the (**Table 2)**.

**Table 1:** List of point mutations identified in the selected transposon mutants.

| Strain | Gene disrupted by transposon insertion | Point mutation identified by whole genome sequencing | Corresponding amino acid substitution |
| --- | --- | --- | --- |
| Tn:35D5 | *bb_0295* | *rpoS* - (T664<sup>A</sup>) | Lysine 222 to stop codon |
| Tn:36C12 | *bb_0421* | *rpoS* - (T664<sup>A</sup>) | Lysine 222 to stop codon |

**Table 2:** List of *B. burgdorferi* strains and plasmids used in the study.

| Strain/plasmid | Description | Source |
| --- | --- | --- |
| 5A14 | Wild-type *B. burgdorferi* | (62) |
| AH130 | Wild-type *B. burgdorferi* | (11) |
| 5A4 | Wild-type *B. burgdorferi* | (63) |
| 5A18NP1 | Wild-type *B. burgdorferi* | (63) |
| $\Delta rpoS$ | *rpoS* mutant in 5A14, 5A4 and 297 background | (6, 8, 64) |
| *rpoS*<sup>com</sup> | *rpoS* complement in 5A14 and 5A4 background | (8, 64) |
| $\Delta rpoN$ | *rpoN* mutant in 5A4NP1 and 297 background | (6, 64) |
| $\Delta bosR$ | *bosR* mutant in 5A14 and 297 background | (18) |
| $\Delta ospC$ | *ospC* mutant in 5A4NP1 | (65) |
| $\Delta bb\_0295$ | *hslU* mutant in 5A18NP1 | Present study |
| $\Delta bb\_0421$ | *bb\_0421* mutant in 5A18NP1 | Present study |
| Tn:001 | Transposon mutant | Present study |
| Tn:002 | Transposon mutant | Present study |
| $\Delta rrp1$ | *rrp1* mutant in 5A4NP1 | Present study |
| *rrp2*<sup>G239C</sup> | *rrp2*<sup>G239C</sup> mutant in 5A4NP1 and 297 background | (11, 12) |
| *rrp2*<sup>G239Ccom</sup> | *rrp2* complement in 5A4NP1 and 297 background | (11, 12) |
| BbSR035 | Wild-type *B. burgdorferi* 5A18NP1 expressing pSR027 | Present study |
| BbSR036 | Wild-type *B. burgdorferi* 5A18NP1 expressing pSR028 | Present study |
| BbSR105 | Wild-type *B. burgdorferi* B31M expressing pOY463 | Present study |
| BbSR151 | *rpoS* mutant in 5A14 expressing *plac*-5'UTR<sub>lac</sub>-*rpoS* shuttle plasmid | (18) |
| BbSR186 | *bosR* mutant in AH130 expressing *plac*-5'UTR<sub>lac</sub>-*bosR* shuttle plasmid | Present study |
| BbSR220 | *rpoS* mutant in 297 expressing *plac*-5'UTR<sub>lac</sub>-*bosR* shuttle plasmid | Present study |
| BbSR221 | *rpoN* mutant in 297 expressing *plac*-5'UTR<sub>lac</sub>-*bosR* shuttle plasmid | Present study |
| BbSR226 | *rpoS* mutant in 297 expressing *plac*-5'UTR<sub>flaB</sub>-*rpoS* shuttle plasmid | Present study |
| BbSR236 | *rpoS* mutant in 5A18NP1 background | Present study |
| BbSR244 | *rpoS* complement in 5A18NP1 background | Present study |
| $\Delta rpoS$ | *rpoS* mutant in 5A18, 5A11, 5A18NP1 and 5A4NP1 background | Present study |
| pSR027 | Suicidal plasmid for the cis complementation of *rpoS* | (18) |
| pSR028 | Suicidal plasmid for creating <sup>T664</sup>A mutation in the chromosomal copy of *rpoS* | Present study |
| pSR069 | Shuttle plasmid containing *plac*-5'UTR*rpoS*-*rpoS* with a *Strep*<sup>R</sup> marker | (18) |
| pSR083 | Shuttle plasmid containing *plac*-5'UTR<sub>lac</sub>-*bosR* with a *Strep*<sup>R</sup> marker | Present study |
| pSR115 | Shuttle plasmid containing *plac*-5'UTR<sub>flaB</sub>-*rpoS* with a *Strep*<sup>R</sup> marker | (18) |
| pOY110 | Shuttle plasmid containing *plac-5'UTR<sub>lac</sub>-rpoS* with a *Strep<sup>R</sup>* marker | (18) |
| pGKT | Suicidal plasmid containing *Himar* transposon | (29) |
| pJK002 | Shuttle plasmid containing *rpoSp-luc* | (18) |
| pOY463 | Shuttle plasmid containing *bosRp-luc* | (36) |

### Generating a genome wide transposon library in *B. burgdorferi*

Random transposon mutagenesis was performed using the infectious *B. burgdorferi* B31 clone 5A18NP1. Briefly, electrocompetent *B. burgdorferi* cells were freshly prepared and were transformed with 5 µg of each plasmid (pGKT) by electroporation following previously published protocols (29, 58). Cultures were allowed to recover overnight in BSK-II medium without antibiotics, followed by limiting dilution and seeding into 96-well plates with 200 µg/ml kanamycin and 50 µg/ml gentamicin. After two weeks of incubation, positive colonies were selected and cultured in liquid BSK-II medium with the same antibiotics until mid-log phase. Cultures were then mixed with equal volume of BSK-II medium containing 30% (v/v) glycerol and were stored at -70°C. The transposon insertion site was determined by rescuing the circularized Hind*III* digested fragments in *E.coli* f (29), with the exact transposon insertion site of each clone in the library was determined by dideoxynucleotide sequencing from recovered plasmid using the specific sequencing primers. Identification of the insertion site was accomplished using batch local BLAST analysis (Bioedit; http://bioedit.software.informer.com).

### Constructing *B. burgdorferi* expressing a K222* mutation RpoS

In order to make a *B. burgdorferi* strain expressing a K222* mutation in the chromosomal copy of RpoS, we utilized the previously developed suicide plasmid pSR027 which carries a wild-type *rpoS* linked to an *aadA* marker (conferring streptomycin resistance) for cis-complementation of the *rpoS* mutant (18). Briefly, employing site-directed mutagenesis approach using Q5® Site-Directed Mutagenesis Kit (New England Biolabs), T to A point mutation was introduced at the 664 nucleotides of *rpoS* ORF in pSR028 using specific sets of primer pSR028 FP: TTAAATTAGTATCTTTCCTTTTCATTTAATTTTG and pSR028 RP: AAAAGATATAACCTGGACAATAGTCC respectively. All the procedure including designing of primers, PCR based mutations followed by KLD (kinase, ligase and DpnI) treatment were performed as per the manufacturer’s guidelines. Mutations were confirmed by sequencing. The resulted suicidal plasmid pSR028 was transformed into wild-type *B. burgdorferi* 5A18NP1 competent cells and transformants were selected based on streptomycin resistance (50 μg/ml).

### Constructing *B. burgdorferi* expressing *lacp*-UTR*_lac_*-*bosR* shuttle plasmid

To artificially induce *bosR* expression in *B. burgdorferi*, an IPTG-inducible *bosR* expression construct was constructed using a *lacp*-based inducible expression system (59). The *bosR* open reding frame was amplified from *B. burgdorferi* genomic DNA using primers pSR083 FP (GATACATATGAACGACAACATAATAGACGTACATTC) and pSR083 RP (GATAAGATCTTCATAAAGTGATTTCCTTGTTCTC). The purified PCR product was digested with NdeI and BglII restriction enzymes and cloned downstream of an T5 promoter into the shuttle plasmid pOY99.2 (17). The resulted shuttle vector pSR083, was transformed into *B. burgdorferi* 297 isogenic mutants of *bosR*, *rpoN* and *rpoS* with transformants selected based on streptomycin resistance (50 μg/ml).

### Immunoblot analysis

Spirochetes from mid-log or stationary phase-grown cultures were harvested by centrifuging at 8000 × g for 10 min, followed by three washes with PBS (pH 7.4) at 4°C. Pellets were suspended in SDS buffer containing 50 mM Tris–HCl (pH 8.0), 0.3% sodium dodecyl sulfate (SDS) and 10 mM dithiothreitol (DTT). Cell lysates (10^8^ cells per lane) were separated by 12% SDS-polyacrylamide gel electrophoresis (PAGE) and transferred to nitrocellulose membranes (GE-Healthcare, Milwaukee, WI). Membranes were probed with mouse monoclonal antibody of anti- BosR (1:3000 dilution), anti-FlaB (1:3000 dilution) or anti-RpoS (1:100 dilution) (16, 60, 61), followed by anti-mouse IgG-HRP secondary antibody (1:1000; Santa Cruz Biotechnology). Horseradish peroxidase activity was detected using enhanced chemiluminescence method (Thermo Pierce ECL Western Blotting Substrate) with subsequent exposure to X-ray film.

### Quantitative real time (qRT-PCR) PCR analyses

RNA samples were extracted from *B. burgdorferi* cultures using the RNeasy mini kit (Qiagen, Valencia, CA) according to the manufacturer’s protocols (60), followed by on-column treatment with RNase-free DNase I treatment Promega, (Madison, WI). The quality of DNA-free RNA was confirmed by PCR amplification of *flaB* of *B. burgdorferi*. cDNA synthesis was performed using the SuperScript III reverse transcriptase with random primers (Invitrogen, Carlsbad, CA). The primers for *bosR* (*bosR* q-RT PCR FP: AGCTTGGCTTCCACAATAGC;*bosR* q-RT PCR RP: TTGCAATGCCCTGAGTAAATGA) were designed using Primer BLAST software. The cycling conditions were as follows: initial denaturation of 94°C for 5 min followed by 35 cycles of denaturation at 94°C for 30 s, primer annealing at 59°C for 30 s, and extension at 72°C for 40 s, with a final melt curve analysis. All reactions were carried out in triplicates using an QuantStudio™ 3 Real-Time PCR thermocycler and were analyzed using QuantStudio™ 3 Real-Time PCR software. Relative transcript levels were normalized to flaB transcript levels, as described previously (60).

### Protein turnover assay

Protein turnover was assessed as previously described (35) using wild-type 5A14, an *rpoS* mutant (with a kanamycin-resistant marker insertion) and the *rpoS*-complemented strain constructed in the same background. Briefly 10^4^ cells/ml of *B. burgdorferi* were inoculated into 60 ml of BSK-II medium, pH 7.5, and cultivated at 37°C to stationary phase (10^8^ cells/ml). Protein synthesis was arrested by adding 100 μg/ml of spectinomycin and 10 ml of cells were harvested at 0, 6, 12 and 24-hours post- arrest for SDS-PAGE followed by immunoblotting for BosR, RpoS and FlaB.

### Cultivation of *B. burgdorferi* within dialysis membrane chambers (DMCs)

Dialysis membrane chambers (DMCs) containing 1×10^3^ organisms diluted from a mid- logarithmic growth culture at 37°C *in vitro*, were implanted into the peritoneal cavities of female Sprague-Dawley rats as previously described (8, 32). The DMCs were recovered on day 13 post-implantation and spirochetes then were harvested, washed with PBS buffer, and then examined by SDS-PAGE, silver staining, and Western blotting analyses.

## DATA AVAILABILITY

All data in this study has been included in the main text.

## ACKNOWLEDGEMENTS

We express our gratitude to Dr. Zhiming Ouyang for generously supplying the strains and plasmids utilized in this study. Special thanks to Dr Youyun Yang and Joleyn Khoo for their invaluable technical assistance in constructing the Tn transposon library. Funding for this research was partly supported by NIH grants AI083640 and AI152235 (to XFY). Additionally, we acknowledge the use of facilities supported by the research facilities improvement program grant number C06 RR015481-01 from the National Center for Research Resources, NIH.

## Abbreviations

Bb: Borreliella burgdorferi
BosR: Borrelia oxidative stress regulatory protein
σ^54^: Sigma 54
σ^S^: Sigma S

## References

1. Steere AC, Strle F, Wormser GP, Hu LT, Branda JA, Hovius JWR, Li X, Mead PS. 2016. Lyme borreliosis. Nat Rev Dis Primers 2:16090–16090.

2. Radolf JD, Caimano MJ, Stevenson B, Hu LT. 2012. Of ticks, mice and men: understanding the dual-host lifestyle of Lyme disease spirochaetes. Nat Rev Micro 10:87–99.

3. Stevenson B, Seshu J. 2017. Regulation of Gene and Protein Expression in the Lyme Disease Spirochete. Curr Top Microbiol Immunol doi:10.1007/82_2017_49.

4. Samuels DS, Lybecker MC, Yang XF, Ouyang Z, Bourret TJ, Boyle WK, Stevenson B, Drecktrah D, Caimano MJ. 2021. Gene Regulation and Transcriptomics. Curr Issues Mol Biol 42:223–266.

5. Stevenson B. 2023. The Lyme disease spirochete, *Borrelia burgdorferi*, as a model vector-borne pathogen: insights on regulation of gene and protein expression. Curr Opin Microbiol 74:102332.

6. Hübner A, Yang X, Nolen DM, Popova TG, Cabello FC, Norgard MV. 2001. Expression of *Borrelia burgdorferi* OspC and DbpA is controlled by a RpoN-RpoS regulatory pathway. Proc Natl Acad Sci U S A 98:12724–9.

7. Caimano MJ, Eggers CH, Gonzalez CA, Radolf JD. 2005. Alternate sigma factor RpoS is required for the in vivo-specific repression of *Borrelia burgdorferi* plasmid lp54-borne *ospA* and *lp6.6* genes. J Bacteriol 187:7845–7852.

8. Caimano MJ, Groshong AM, Belperron A, Mao J, Hawley KL, Luthra A, Graham DE, Earnhart CG, Marconi RT, Bockenstedt LK, Blevins JS, Radolf JD. 2019. The RpoS Gatekeeper in *Borrelia burgdorferi*: An Invariant Regulatory Scheme That Promotes Spirochete Persistence in Reservoir Hosts and Niche Diversity. Front Microbiol 10.

9. Fisher MA, Grimm D, Henion AK, Elias AF, Stewart PE, Rosa PA, Gherardini FC. 2005. *Borrelia burgdorferi* σ^54^ is required for mammalian infection and vector transmission but not for tick colonization. Proc Natl Acad Sci U S A 102:5162–5167.

10. Ouyang Z, Narasimhan S, Neelakanta G, Kumar M, Pal U, Fikrig E, Norgard M. 2012. Activation of the RpoN-RpoS regulatory pathway during the enzootic life cycle of *Borrelia burgdorferi*. BMC Microbiol 12:44.

11. Yang XF, Alani SM, Norgard MV. 2003. The response regulator Rrp2 is essential for the expression of major membrane lipoproteins in *Borrelia burgdorferi*. Proc Natl Acad Sci U S A 100:11001–11006.

12. Boardman BK, He M, Ouyang Z, Xu H, Pang X, Yang XF. 2008. Essential role of the response regulator Rrp2 in the infectious cycle of *Borrelia burgdorferi*. Infect Immun 76:3844–3853.

13. Blevins JS, Xu H, He M, Norgard MV, Reitzer L, Yang XF. 2009. Rrp2, a sigma54-dependent transcriptional activator of *Borrelia burgdorferi*, activates *rpoS* in an enhancer-independent manner. J Bacteriol 191:2902–5.

14. Ouyang Z, Zhou J. 2016. The putative Walker A and Walker B motifs of Rrp2 are required for the growth of *Borrelia burgdorferi*. Mol Microbiol.

15. Yin Y, Yang Y, Xiang X, Wang Q, Yang Z-N, Blevins J, Lou Y, Yang XF. 2016. Insight into the dual functions of bacterial enhancer-binding protein Rrp2 of *Borrelia burgdorferi*. J Bacteriol 198:1543–1552.

16. Hyde JA, Shaw DK, Smith Iii R, Trzeciakowski JP, Skare JT. 2009. The BosR regulatory protein of *Borrelia burgdorferi* interfaces with the RpoS regulatory pathway and modulates both the oxidative stress response and pathogenic properties of the Lyme disease spirochete. Mol Microbiol 74:1344–55.

17. Ouyang Z, Deka RK, Norgard MV. 2011. BosR (BB0647) controls the RpoN-RpoS regulatory pathway and virulence expression in Borrelia burgdorferi by a novel DNA-binding mechanism. PLoS Pathog 7:e1001272.

18. Raghunandanan S, Priya R, Alanazi F, Lybecker MC, Schlax PJ, Yang XF. 2024. A Fur family protein BosR is a novel RNA-binding protein that controls *rpoS* RNA stability in the Lyme disease pathogen. Nucleic Acids Res:gkae114.

19. Schwan TG, Piesman J, Golde WT, Dolan MC, Rosa PA. 1995. Induction of an outer surface protein on *Borrelia burgdorferi* during tick feeding. Proceedings of the National Academy of Sciences USA 92:2909–2913.

20. Yang X, Goldberg MS, Popova TG, Schoeler GB, Wikel SK, Hagman KE, Norgard MV. 2000. Interdependence of environmental factors influencing reciprocal patterns of gene expression in virulent *Borrelia burgdorferi*. Mol Microbiol 37:1470–1479.

21. Carroll JA, Cordova RM, Garon CF. 2000. Identification of 11 pH-regulated genes in *Borrelia burgdorferi* localizing to linear plasmids. Infection and Immunity 68:6677–6684.

22. Seshu J, Boylan JA, Gherardini FC, Skare JT. 2004. Dissolved oxygen levels alter gene expression and antigen profiles in Borrelia burgdorferi. Infection and Immunity 72:1580–1586.

23. Hyde JA, Trzeciakowski JP, Skare JT. 2007. *Borrelia burgdorferi* alters its gene expression and antigenic profile in response to CO_2_ levels. J Bacteriol 189:437–445.

24. Troxell B, Ye M, Yang Y, Carrasco SE, Lou Y, Yang XF. 2013. Manganese and Zinc Regulate Virulence Determinants in *Borrelia burgdorferi*. Infect Immun 81:2743–2752.

25. Caimano MJ, Iyer R, Eggers CH, Gonzalez C, Morton EA, Gilbert MA, Schwartz I, Radolf JD. 2007. Analysis of the RpoS regulon in Borrelia burgdorferi in response to mammalian host signals provides insight into RpoS function during the enzootic cycle. Mol Microbiol 65:1193–1217.

26. Lin Y-H, Chen Y, Smith TC, Karna SLR, Seshu J. 2018. Short-Chain Fatty Acids Alter Metabolic and Virulence Attributes of *Borrelia burgdorferi*. Infect Immun 86:e00217–18.

27. Xu H, Caimano MJ, Lin T, He M, Radolf JD, Norris SJ, Gheradini F, Wolfe AJ, Yang XF. 2010. Role of acetyl-phosphate in activation of the Rrp2-RpoN-RpoS pathway in *Borrelia burgdorferi*. PLoS Pathog 6:e1001104. doi: 10.1371/journal.ppat.1001104

28. Stevenson B, Schwan TG, Rosa PA. 1995. Temperature-related differential expression of antigens in the Lyme disease spirochete, Borrelia burgdorferi. Infect Immunity 63:4535–4539.

29. Stewart PE, Hoff J, Fischer E, Krum JG, Rosa PA. 2004. Genome-wide transposon mutagenesis of *Borrelia burgdorferi* for identification of phenotypic mutants. Appl Environ Microbiol 70:5973–5979.

30. Lin T, Gao L, Zhang C, Odeh E, Jacobs MB, Coutte L, Chaconas G, Philipp MT, Norris SJ. 2012. Analysis of an Ordered, Comprehensive STM Mutant Library in Infectious Borrelia burgdorferi Insights into the Genes Required for Mouse Infectivity. PLoS ONE 7:e47532.

31. Mittal N, Roy N, Babu MM, Janga SC. 2009. Dissecting the expression dynamics of RNA-binding proteins in posttranscriptional regulatory networks. Proc Natl Acad Sci U S A 106:20300–20305.

32. Akins DR, Bourell KW, Caimano MJ, Norgard MV, Radolf JD. 1998. A new animal model for studying Lyme disease spirochetes in a mammalian host-adapted state. J Clin Invest 101:2240–2250.

33. Duval M, Simonetti A, Caldelari I, Marzi S. 2015. Multiple ways to regulate translation initiation in bacteria: mechanisms, regulatory circuits, dynamics. Biochimie 114:18–29.

34. Borovinskaya MA, Shoji S, Holton JM, Fredrick K, Cate JH. 2007. A steric block in translation caused by the antibiotic spectinomycin. ACS Chem Biol 2:545–552.

35. Sze CW, Zhang K, Lynch MJ, Iyer R, Crane BR, Schwartz I, Li C. 2023. A chemosensory-like histidine kinase is dispensable for chemotaxis in vitro but regulates the virulence of *Borrelia burgdorferi* through modulating the stability of RpoS. PLoS Pathog 19:e1011752.

36. Ouyang Z, Zhou J, Norgard MV. 2016. Evidence that BosR (BB0647) is a positive autoregulator in *Borrelia burgdorferi*. Infect Immun 84:2566–2574.

37. Mahmoud SA, Chien P. 2018. Regulated proteolysis in bacteria. Annu Rev Biochem 87:677–696.

38. He M, Zhang J-J, Ye M, Lou Y, Yang XF. 2014. The cyclic di-GMP receptor PlzA controls virulence gene expression through RpoS in *Borrelia burgdorferi*. Infect Immun 82:445–52.

39. Wang P, Yu Z, Santangelo TJ, Olesik J, Wang Y, Heldwein E, Li X. 2017. BosR Is A Novel Fur Family Member Responsive to Copper and Regulating Copper Homeostasis in *Borrelia burgdorferi*. J Bacteriol 199.

40. Saputra EP, Trzeciakowski JP, Hyde JA. 2020. *Borrelia burgdorferi* spatiotemporal regulation of transcriptional regulator *bosR* and decorin binding protein during murine infection. Sci Rep 10:12534.

41. Medrano MS, Ding Y, Wang X-G, Lu P, Coburn J, Hu LT. 2007. Regulators of expression of the oligopeptide permease A proteins of *Borrelia burgdorferi*. J Bacteriol 189:2653–2659.

42. Muffler A, Fischer D, Altuvia S, Storz G, Hengge-Aronis R. 1996. The response regulator RssB controls stability of the sigma (S) subunit of RNA polymerase in Escherichia coli. EMBO J 15:1333–1339.

43. Studemann A, Noirclerc-Savoye M, Klauck E, Becker G, Schneider D, Hengge R. 2003. Sequential recognition of two distinct sites in sigma(S) by the proteolytic targeting factor RssB and ClpX. EMBO J 22:4111–20.

44. Zhou Y, Gottesman S, Hoskins JR, Maurizi MR, Wickner S. 2001. The RssB response regulator directly targets ςS for degradation by ClpXP. Genes Dev 15:627–637.

45. Wölflingseder M, Tutz S, Fengler VH, Schild S, Reidl J. 2022. Regulatory interplay of RpoS and RssB controls motility and colonization in Vibrio cholerae. Int J Med Microbiol 312:151555.

46. Bouillet S, Hamdallah I, Majdalani N, Tripathi A, Gottesman S. 2024. A negative feedback loop is critical for recovery of RpoS after stress in *Escherichia coli*. Plos Genet 20:e1011059.

47. Ojaimi C, Brooks C, Casjens S, Rosa P, Elias A, Barbour A, Jasinskas A, Benach J, Katona L, Radolf J, Caimano M, Skare J, Swingle K, Akins D, Schwartz I. 2003. Profiling of temperature-induced changes in *Borrelia burgdorferi* gene expression by using whole genome arrays. Infect Immun 71:1689–1705.

48. Arnold WK, Savage CR, Brissette CA, Seshu J, Livny J, Stevenson B. 2016. RNA-Seq of *Borrelia burgdorferi* in Multiple Phases of Growth Reveals Insights into the Dynamics of Gene Expression, Transcriptome Architecture, and Noncoding RNAs. PLoS ONE 11:e0164165.

49. Sze CW, Smith A, Choi YH, Yang X, Pal U, Yu A, Li C. 2013. Study of the response regulator Rrp1 reveals its regulatory role in chitobiose utilization and virulence of *Borrelia burgdorferi*. Infect Immun 81:1775–1787.

50. Drecktrah D, Lybecker M, Popitsch N, Rescheneder P, Hall LS, Samuels DS. 2015. The *Borrelia burgdorferi* RelA/SpoT homolog and stringent response regulate survival in the tick vector and global gene expression during starvation. PLoS Pathog 11:e1005160.

51. Miller CL, Karna SLR, Seshu J. 2013. *Borrelia* host adaptation regulator (BadR) regulates *rpoS* to modulate host adaptation and virulence factors in *Borrelia burgdorferi*. Mol Microbiol 88:105–24.

52. Ouyang Z, Zhou J. 2015. BadR (BB0693) controls growth phase-dependent induction of *rpoS* and *bosR* in *Borrelia burgdorferi* via recognizing TAAAATAT motifs. Mol Microbiol 98:1147–1167.

53. Grassmann AA, Tokarz R, Golino C, McLain MA, Groshong AM, Radolf JD, Caimano MJ. 2023. BosR and PlzA reciprocally regulate RpoS function to sustain *Borrelia burgdorferi* in ticks and mammals. J Clin Investig 133.

54. Shi Y, Dadhwal P, Li X, Liang FT. 2014. BosR functions as a repressor of the *ospAB* operon in *Borrelia burgdorferi*. PLoS One 9:e109307.

55. Wang P, Dadhwal P, Cheng Z, Zianni MR, Rikihisa Y, Liang FT, Li X. 2013. *Borrelia burgdorferi* oxidative stress regulator BosR directly represses lipoproteins primarily expressed in the tick during mammalian infection. Mol Microbiol 89:1140–1153.

56. Grove AP, Liveris D, Iyer R, Petzke M, Rudman J, Caimano MJ, Radolf JD, Schwartz I. 2017. Two Distinct Mechanisms Govern RpoS-Mediated Repression of Tick-Phase Genes during Mammalian Host Adaptation by *Borrelia burgdorferi*, the Lyme Disease Spirochete. mBio 8.

57. Barbour AG. 1984. Isolation and cultivation of Lyme disease spirochetes. Yale J Biol Med 57:521–5.

58. Samuels DS. 1995. Electrotransformation of the spirochete Borrelia burgdorferi. Electroporation protocols for microorganisms:253–259.

59. Blevins JS, Revel AT, Smith AH, Bachlani GN, Norgard MV. 2007. Adaptation of a luciferase gene reporter and *lac* expression system to *Borrelia burgdorferi*. Appl Environ Microbiol 73:1501–1513.

60. Zhang Y, Chen T, Raghunandanan S, Xiang X, Yang J, Liu Q, Edmondson DG, Norris SJ, Yang XF, Lou Y. 2020. YebC regulates variable surface antigen VlsE expression and is required for host immune evasion in *Borrelia burgdorferi*. PLoS Pathog 16:e1008953.

61. Zhang J-J, Raghunandanan S, Wang Q, Priya R, Alanazi F, Lou Y, Yang XF. 2024. BadR directly represses the expression of the glycerol utilization operon in the Lyme disease pathogen. J Bacteriol:e00340–23.

62. Purser JE, Norris SJ. 2000. Correlation between plasmid content and infectivity in Borrelia burgdorferi Proc Natl Acad Sci U S A 97:13865–13870.

63. Kawabata H, Norris SJ, Watanabe H. 2004. BBE02 disruption mutants of *Borrelia burgdorferi* B31 have a highly transformable, infectious phenotype. Infect Immun 72:7147–7154.

64. Alanazi F, Raghunandanan S, Priya R, Yang XF. 2023. The Rrp2-RpoN-RpoS pathway plays an important role in the blood-brain barrier transmigration of the Lyme disease pathogen. Infect Immun 91:e00227–23.

65. He M, Oman T, Xu H, Blevins J, Norgard MV, Yang XF. 2008. Abrogation of *ospAB* constitutively activates the Rrp2-RpoN-RpoS pathway (sigmaN-sigmaS cascade) in *Borrelia burgdorferi*. Mol Microbiol 70:1453–64.

